# Automated joint skull-stripping and segmentation with Multi-Task U-Net in large mouse brain MRI databases

**DOI:** 10.1101/2020.02.25.964015

**Authors:** Riccardo De Feo, Artem Shatillo, Alejandra Sierra, Juan Miguel Valverde, Olli Gröhn, Federico Giove, Jussi Tohka

## Abstract

Skull-stripping and region segmentation are fundamental steps in preclinical magnetic resonance imaging (MRI) studies, and these common procedures are usually performed manually. We present Multi-task U-Net (MU-Net), a convolutional neural network designed to accomplish both tasks simultaneously. MU-Net achieved higher segmentation accuracy than state-of-the-art multi-atlas segmentation methods with an inference time of 0.35 seconds and no pre-processing requirements. We evaluated the performance of our network in the presence of skip connections and recently proposed framing connections, finding the simplest network to be the most effective. We tested MU-Net with an unusually large dataset combining several independent studies consisting of 1,782 mouse brain MRI volumes of both healthy and Huntington animals, and measured average Dice scores of 0.906 (striati), 0.937 (cortex), and 0.978 (brain mask). These high evaluation scores demonstrate that MU-Net is a powerful tool for segmentation and skull-stripping, decreasing inter and intra-rater variability of manual segmentation. The MU-Net code and the trained model are publicly available at https://github.com/Hierakonpolis/MU-Net.

## 1 Introduction

Preclinical imaging studies serve a fundamental role in biological and medical research, relating research results at the molecular level to clinical application in diagnosis and therapy. Magnetic Resonance Imaging (MRI) represents approximately 23% of all small-animal imaging studies providing the opportunity to monitor the development of pathological conditions and responses to treatment in a non-invasive way [1]. Its unique qualities also include the availability of different imaging contrasts, rendering MRI extremely useful in the context of preclinical neuroscience with applications from drug development [2] to basic research [3].

Skull-stripping and region segmentation represent an integral part of processing pipelines in murine MR imaging [4, 5]. Skull-stripping refers to the identification of the brain within the MRI volume and region segmentation refers to the labeling of specific anatomical regions of interest (ROIs) within the brain. In preclinical MRI, these tasks are often performed manually. While manual segmentation represents the gold standard and is employed as the ground truth when evaluating automated segmentation algorithms, it is time-consuming and depends on the expertise of the annotators performing the segmentation. Furthermore, manual segmentation suffers from both intra- and inter-rater variability, both in small animal [6] and human MRI [7, 8].

In preclinical MRI, state-of-the-art automated region segmentation pipelines are based on atlas registration: individual MRI volumes are aligned with a labeled template (atlas) and the labels propagated to the individual volumes [9–13]. The accuracy of registration-based segmentation depends both on the suitability of the template and the choice of the registration algorithm. The segmentation accuracy can be improved by multi-atlas strategies, where multiple atlases are registered to the same volume and so-resulting segmentation maps are combined via, for example, majority voting. Regarding multi-atlas strategies in mouse MRI, Bai et al. [14] compared different single and multi-atlas methods for atlas-based segmentation of the mouse brain finding that the combination of a diffeomorphic registration algorithm and multi-atlas segmentation provided the most accurate results. Ma et al. [15] demonstrated that the multi-atlas methods are superior to single-atlas methods and the STEPS procedure for combining segmentations [16] brings advantages over earlier combination methodologies. While multi-atlas segmentation accounts for individual variability more effectively than single-atlas segmentation, it also requires multiple labeled atlases and multiple registration steps, significantly increasing the segmentation time.

Instead of directly employing one or more manually segmented atlases, deep neural networks (DNNs) [17] can use these as training data to learn a mapping function from the image data to the segmentation maps. In this way, the anatomical information is not explicitly represented in a set of maps but implicitly encoded in the trained network. DNNs, and in particular Convolutional Neural Networks (CNNs), have been successfully applied in a large number of computer vision tasks, including segmentation of human brain MRI. For example, Wachinger et al. [18] developed a region segmentation CNN significantly outperforming state-of-the-art, registration-based methods for the healthy human brain, both in terms of inference time and accuracy. Roy et al. [19] further improved on both aspects with a network based on the U-Net architecture [20], with a reported segmentation time of 20 s for a brain scan. However, within small-animal MRI, the applications of CNNs have been limited to skull-stripping: Roy et al. [21] trained a Convolutional Neural Network (CNN) algorithm based on Google Inception [22] for the skull-stripping in humans and mice after traumatic brain injury, achieving better performance than other state-of-the-art methods (3D Pulse Coupled Neural Networks (3D-PCNN) [23] and Rapid Automatic Tissue Segmentation (RATS) [24]).

A specific type of CNN architecture, U-Net, has proved to be valuable in biomedical image segmentation. U-Net is based on the encoder/decoder structure, adding skip connections between the encoder branch and the decoder branch, allowing it to easily integrate multi-scale information and better propagate the gradient during training. This architecture has been shown to generalize even from a limited amount of annotated data [25], and as such is well suited for medical imaging, where datasets as large as the ones commonly used for CNNs are rare. Valverde et al. [26] recently demonstrated the effectiveness of U-Net-like architectures in preclinical research, designing the first DNN for the segmentation of ischemic lesions in rodents and achieving segmentation accuracy comparable or better to inter-rater agreement in manual segmentation.

In this work, we introduce multi-task U-Net (MU-Net) to simultaneously perform skull-stripping and region segmentation of the mouse brain, based on the U-Net architecture. Our main training data consisted of 128 T_2_ MRI volumes from 32 mice at 4 different ages as well as five manually annotated regions (cortex, hippocampi, ventricles, striati and brain mask) from these images. This dataset represents MR images typically employed in drug development. We tested MU-Net in an independent test set of 1782 MRI volumes acquired over the course of four years from both wild type (WT) and Huntington (HT) C57BL/6J mice, allowing us to evaluate MU-Net in a variety of experimental conditions. We demonstrate that with these data MU-Net achieves a significantly higher accuracy than state-of-the-art multi-atlas segmentation methods [16] with a fraction of a segmentation time (approximately 0.35 s). Additionally, we trained MU-Net for the segmentation of mouse MRI with isotropic voxels into 37 ROIs and demonstrate that the segmentation accuracy of MU-Net was equal or better than a state-of-the-art multi-atlas segmentation method [15].

## 2 Results

### 2.1 MU-Net

MU-Net is a U-Net-based network combining region segmentation and skull-stripping to build a shared representation for both tasks, providing at the same time a more efficient feature encoding and a regularizing effect. We present an overview of MU-Net’s architecture in Figure 1. Briefly, MU-Net presents an encoder-decoder U-Net-like architecture, with each branch articulated in four dense blocks. Unlike U-Net, the final block of the decoder branch further branches into two different output maps representing our two tasks, sharing the same feature representation. MU-Net was trained using the Adam optimizer [27] to jointly minimize the Dice loss for both skull-stripping and region segmentation [28]. For further details refer to section 4.4.

**Fig 1.**
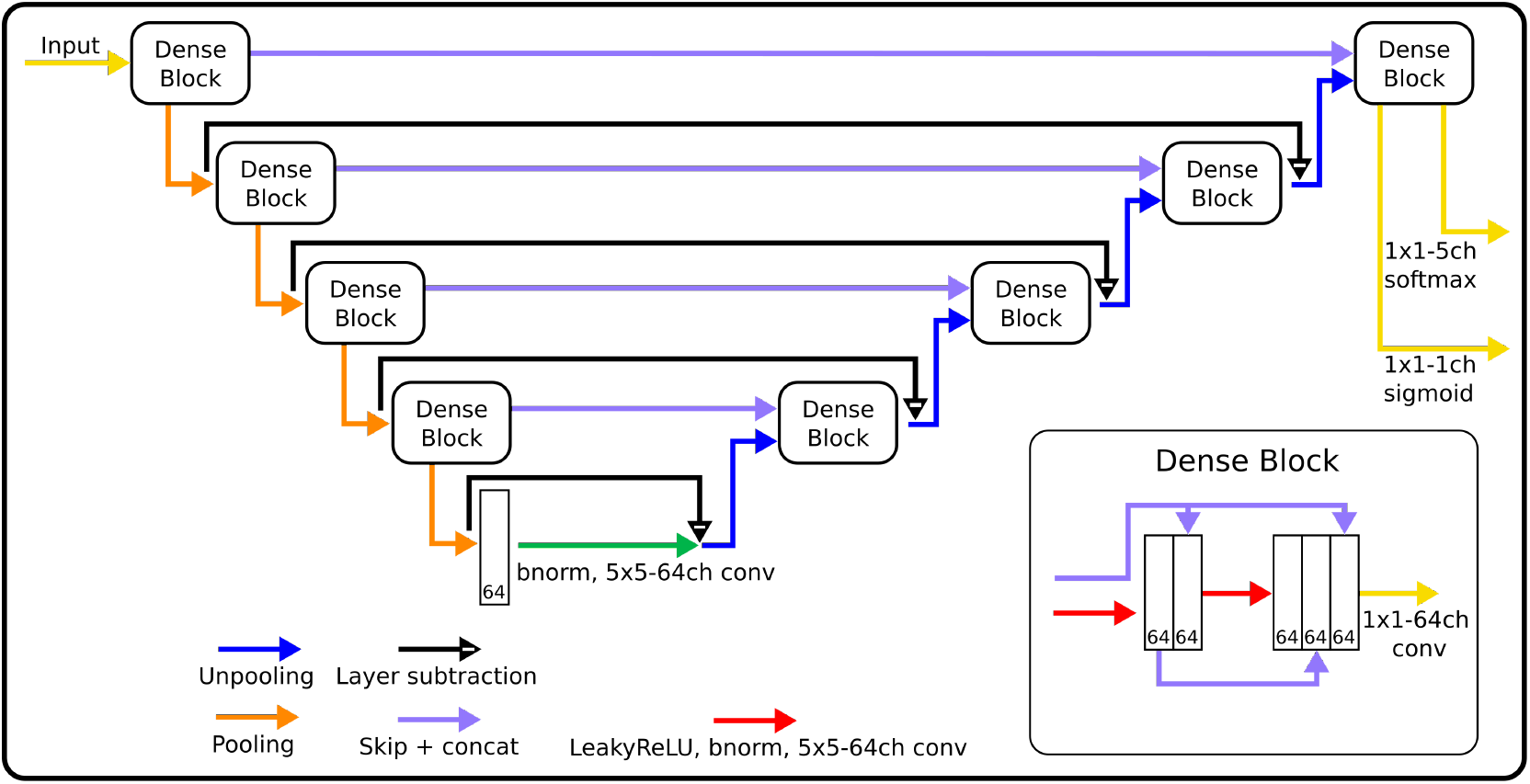
General outline of the architectural features implemented and compared in the networks discussed, varying according to the presence or absence of the in-block dense connections (purple arrows in the dense block box), presence or absence of the layer subtraction connections (black), and the use of 2D or 3D filters.

MU-Net was trained and validated on 128 mouse MRI volumes acquired from 32 animals at four different ages (5, 12, 16, and 32 weeks). The training set was annotated for the brain mask and four more regions: cortex, striati, hippocampi and ventricles, each category including both left and right regions. For validation, we used 5-folds cross validation and measured overlaps using the Dice score [29]. The hypothesis tests between the Dice coefficients of different segmentation method were performed using permutation tests and all the reported p-values are obtained from a permutation test. Validation and metrics are further expanded upon in section 4.3.

Using the training and validation dataset, we compared the performance of different network architectures based on the presence of skip connections, framing connections and 2D or 3D filters. Furthermore, we compared MU-Net with multi-atlas segmentation and evaluated the impact of mouse age on the accuracy of our segmentation maps. Finally, we tested MU-Net on a large dataset including 1782 MRI volumes from 817 mice. These volumes were acquired as parts of 10 studies on Huntington’s disease and included both WT and HT mice with ages ranging from 4 to 60 weeks. This heterogeneous dataset included three segmented regions, for which we measured average Dice scores of 0.906 (striati), 0.937 (cortex), and 0.978 (brain mask) between MU-Net and manual segmentations.

### 2.2 Architecture comparison

We compared the performance of different networks trained with and without dense connections and dual framing connections, in both 2D and 3D implementations. By a dense connection we refer to skip connections inside each dense block of the MU-Net architecture. Dual framing connections refer to the skip connections originally introduced by Han and Ye [30] for an image reconstruction task.

As shown in Table 1, all MU-Nets achieved Dice scores with the ground truth comparable to or higher than the typical inter-rater variability of manual segmentation in the mouse brain (Dice scores from 0.80 to 0.90 [6]). The skull-stripping task achieved an excellent Dice score of 0.984. The ventricles were characterized by the lowest segmentation performance (average Dice score 0.907), while the cortex displayed the highest overlap with the ground truth (average Dice score 0.966). Dice scores for each animal in all ROIs are provided as a supplementary csv-file.

**Table 1.**
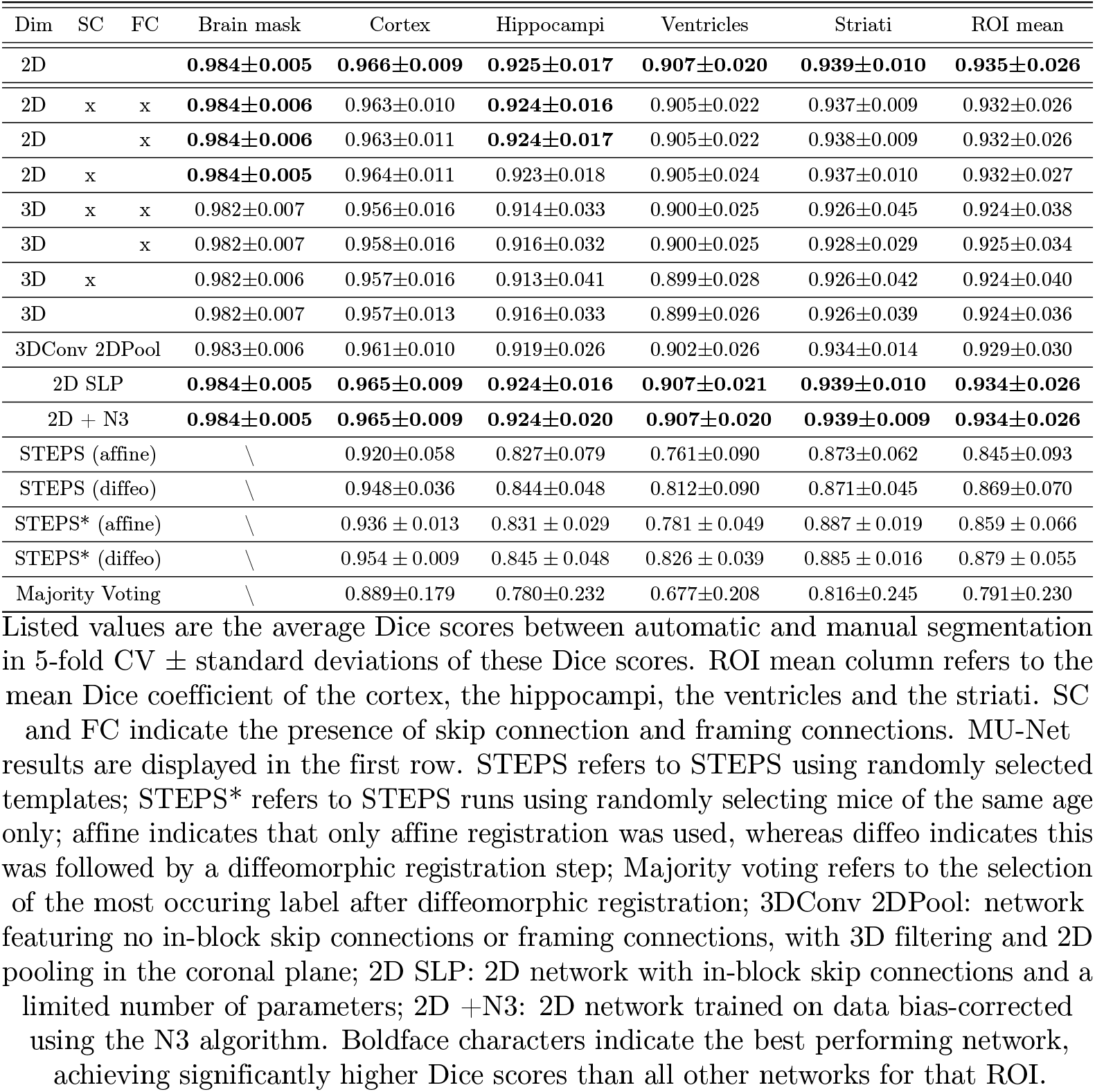
CNN and STEPS accuracies measured using Dice coefficient across different methodological choices.

The network displaying the highest average Dice scores was, in fact, the simplest one, including no in-block skip connections nor framing connections, and using 2D convolutions. The accuracy of this network was significantly higher than the accuracy of other all other 2D networks (*p* < 0.00003, all the hypothesis testing in this work was performed by a paired permutation test as detailed in Section 4.3). Because of its excellent performance and simplicity this network is our choice for the MU-Net architecture.

The choice between 2D and 3D architectures was the most important factor in increasing performance, resulting in a marked increase in mean Dice scores for both tasks (*p* < 0.00001) between all 2D networks compared to the 3D ones. We further compared MU-Net with one featuring less channels per filter (49, 49, 50, 50, from the shallowest to the deepest dense block) to match the number of parameters to the number of parameters of the simplest 2D network. We registered a slightly (but not significantly, *p* = 0.077) lower accuracy compared to MU-Net, indicated as 2D SLP in Table 1.

To test whether the increased performance of 2D architectures compared to the 3D implementation depended on the reduced number of parameters or on an excessive loss of information when pooling in the anterior-posterior direction, we trained a network using 3D filters while only pooling in the coronal plane. This network achieved intermediate performance between the 3D and 2D implementations (Table 1), suggesting that both above mentioned aspects were relevant in increasing the algorithm’s performance.

We trained MU-Net on MR images without bias-correction and on bias-corrected MR images using the N3 method [31]. The validation accuracy achieved with bias correction was indistinguishable from the accuracy of MU-Net trained without bias correction (see Table 1).

### 2.3 Age stratified training sets

We evaluated the performance of MU-Net when restricting the training set to mice of a specific age. Networks trained on data from mice of 12, 16 and 32 weeks achieved higher accuracy, both on their respective validation set and the overall ground truth, compared to the networks trained on 5 weeks mice (*p* < 0.00001). As shown in Fig. 5, while all networks trained on one specific age displayed a statistically significant (*p* < 0.05, unpaired) decrease in mean accuracy when validated on animals of a different age, this difference was highest between the 5 weeks data and the other datasets.

Limiting the training data to one specific age implies that these networks were trained only on a quarter of the data used to train the networks in section 2.2. Irrespective of that, these networks still achieved average Dice score on the mixed-age validation dataset comparable with the accuracy of manual segmentation. The worst performing CNN was the network trained on 5 weeks old mice. Training on the 12, 16 and 32 weeks data and validating on mice of the same age, we observed Dice scores comparable with the overall performance of MU-Net trained on the entire dataset (*p* > 0.15, unpaired). However, we measured a lower overall performance when including mice of all ages in the validation data (*p* < 0.00001), slightly overfitting for each specific age.

### 2.4 Comparison with multi-atlas segmentation

We compared MU-Net with multi-atlas segmentation, applying the state-of-the-art STEPS [16, 32] label fusion method to combine the labels obtained from the registration of multiple labeled volumes. STEPS takes into account local and global image matching, combining an expectation-maximization approach with Markov Random Fields to improve on the segmentation based on the quality of the registration itself. We repeated this procedure using both diffeomorphic and affine registration methods, with randomly-selected templates restricted to same-age mice. The brain mask segmentation was not evaluated as the manually drawn mask was used during the diffeomorphic registration procedure.

MU-Net achieved higher accuracy than all STEPS implementations (*p* < 0.00001). There was a marked qualitative difference between STEPS segmentation and MU-Net (Fig. 2), the latter achieving results visually indistinguishable from manual segmentation.

**Fig 2.**
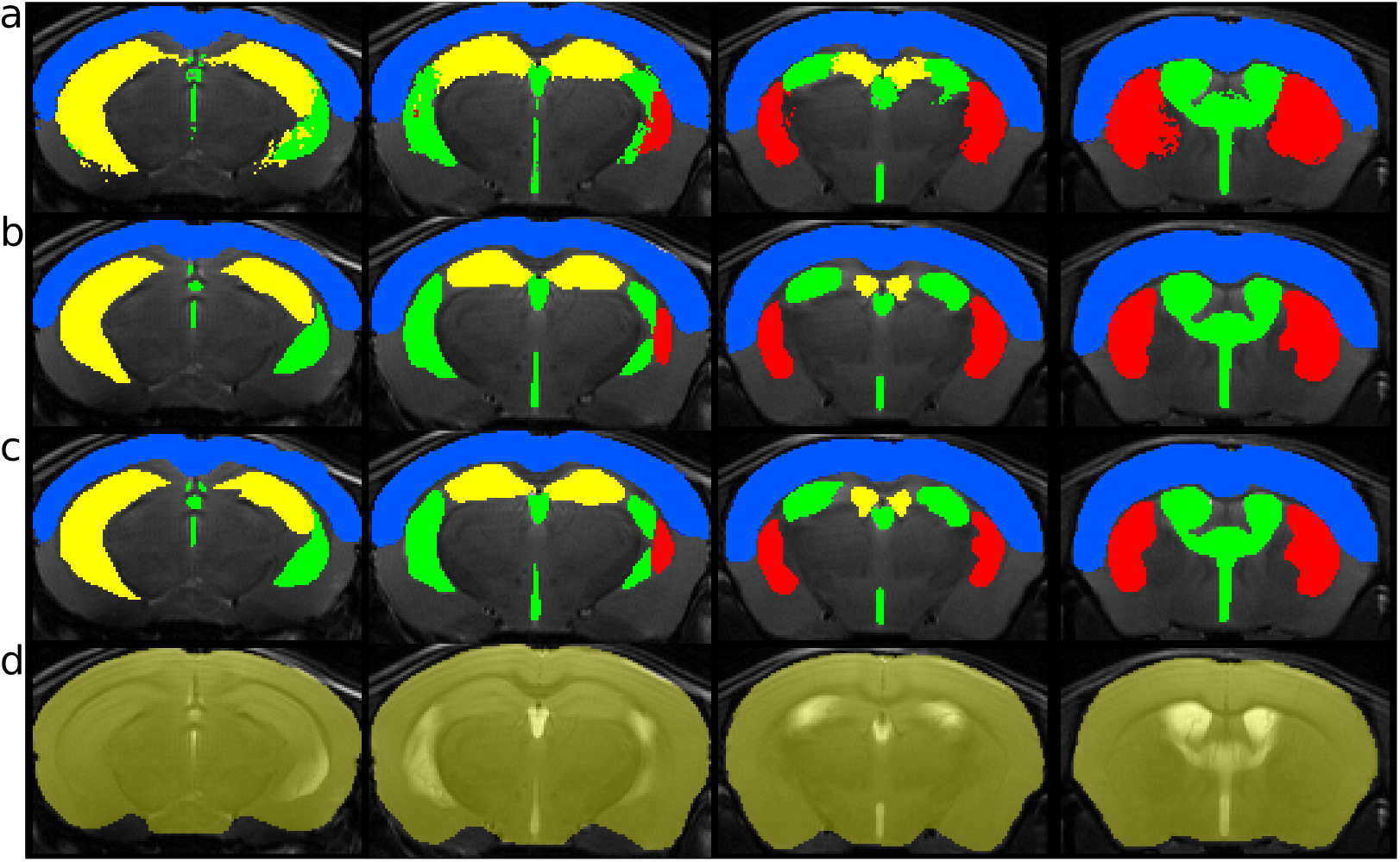
Segmentation comparison in four slices from a single animal: (a) STEPS, (b) MU-Net, and (c) manual annotation. In (a)-(c), the regions highlighted are the cortex (blue), ventricles (green), striati (red), and hippocampi (yellow). Panel (d) shows the inferred brain mask by MU-Net.

MU-Net had an inference time of about 0.35 s (one forward pass) and a training time of 12 hours. STEPS segmentation procedure required total inference time of 117 minutes for each labeled volume (on average 440 s for each pairwise diffeomorphic registration and 7.85 s for label fusion). Implementing STEPS segmentation using only templates of the same age led to a small but significant (*p* < 0.0007) performance improvement over randomly choosing templates of any age. The employment of diffeomorphic registration was the most important factor affecting the performance of STEPS, as displayed in Table 1. A simple majority voting strategy led to significantly lower performance in all ROIs compared to all other label fusion strategies (*p* < 0.003).

Furthermore, we trained MU-Net on the outputs of the implemented STEPS procedures and measured the Dice scores of each network’s output with the ground truth (Table 2). As evidenced in Tables 1 and 2, and Figure 3, MU-Net trained on STEPS segmentations achieved higher Dice score with the ground truth than the same STEPS segmentations constituting the training sets of MU-Net (*p* < 0.00001). With the exception of the network trained on 5 weeks old mice, these hybrid networks were still under-performing compared to training on manually segmented data (*p* < 0.00001).

**Fig 3.**
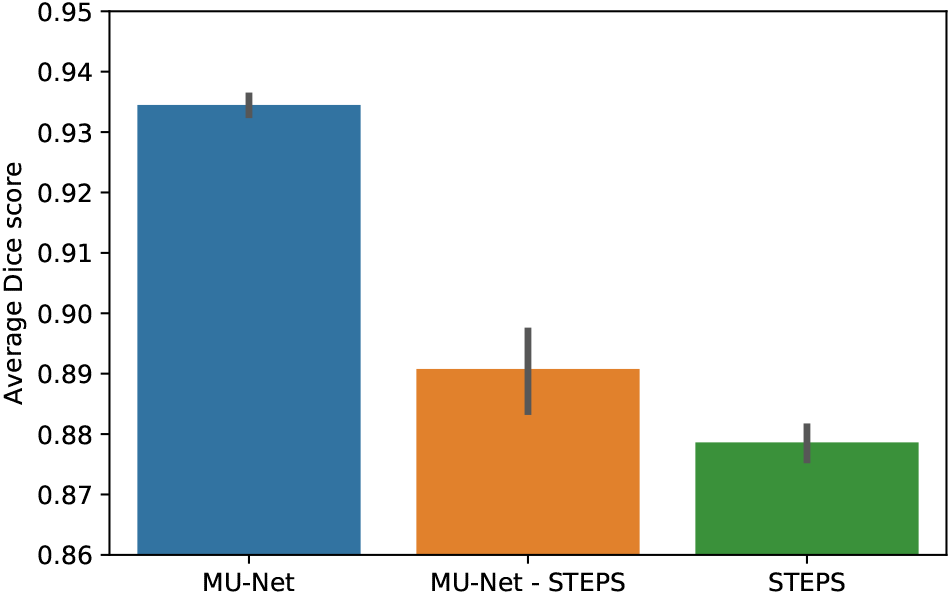
Average Dice score comparison between different segmentation methods, across all ROIs. MU-Net: MU-Net trained on the manually segmented data;MU-Net - STEPS: MU-Net trained on volumes segmented using STEPS; STEPS: STEPS segmentation. The error bar represents standard deviation.

**Table 2.** Mean and standard deviation of average Dice scores evaluating the accuracy of MU-Net trained on volumes segmented via STEPS.

| Training Set | Cortex | Hippocampus | Ventricles | Striatum | ROI mean |
| --- | --- | --- | --- | --- | --- |
| STEPS* | 0.954±0.011 | 0.867±0.027 | 0.866±0.035 | 0.898±0.017 | 0.896±0.043 |
| STEPS | 0.953±0.009 | 0.872±0.022 | 0.849±0.041 | 0.885±0.016 | 0.890±0.046 |

### 2.5 Evaluation on a large number of ROIs with MRM NeAt dataset

We trained and evaluated MU-Net on the MRM NeAt datasets that includes atlases of 10 individual T_2*_-weighted in vivo brain MR images of 12-14 weeks old C57BL/6J mice; each with 37 manually labelled anatomical structures [33]. This same database was selected by Ma et al. [15] to evaluate the STEPS multi-atlas segmentation algorithm on mouse brain MRI. To compare MU-Net with STEPS, we followed the STEPS implementation by Ma et al. [15] as released by the authors.

We used a 5-fold cross validation scheme for evaluation (8 templates for training and 2 templates for testing). As displayed in Fig. 4, the accuracy of MU-Net was comparable or better to STEPS: while in a majority of regions MU-Net’s accuracy was higher than the accuracy of STEPS, this was statistically significant only for the brain mask, external capsule, hypothalamus and brain stem. In the left inferior colliculi, STEPS achieved significantly higher accuracy than MU-Net. Averaging the Dice coefficients across all ROIs, we measured an average Dice score of 0.820 ± 0.031 for MU-Net and 0.814 ± 0.023 for STEPS. While the average accuracy for MU-Net was higher, this difference was not statistically significant (*p* = 0.170). The computation time required by STEPS to segment a single volume was of approximately 20 minutes while MU-Net required less than one second to segment a single volume.

**Fig 4.**
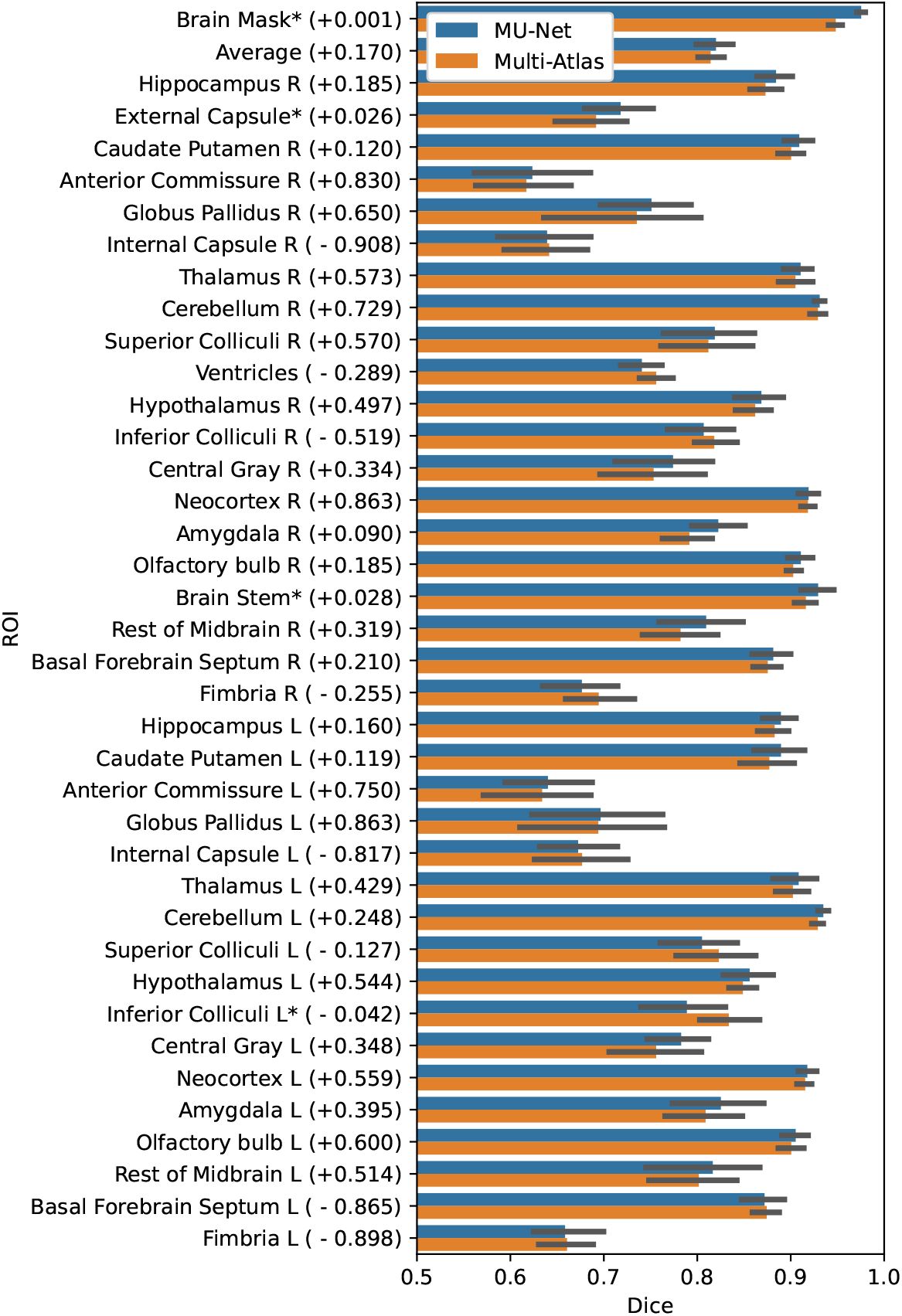
Comparison between the average Dice coefficients of MU-Net and STEPS multi-atlas algorithm by Ma et al. Error bars correspond to standard deviation for the average accuracy. Permutation-test based p-values for each comparison are provided in parentheses after the ROI name, + indicates that the average Dice coefficient for MU-Net was higher and indicates that the average Dice coefficient for STEPS was higher, * indicates a statistically significant difference.

**Fig 5.**
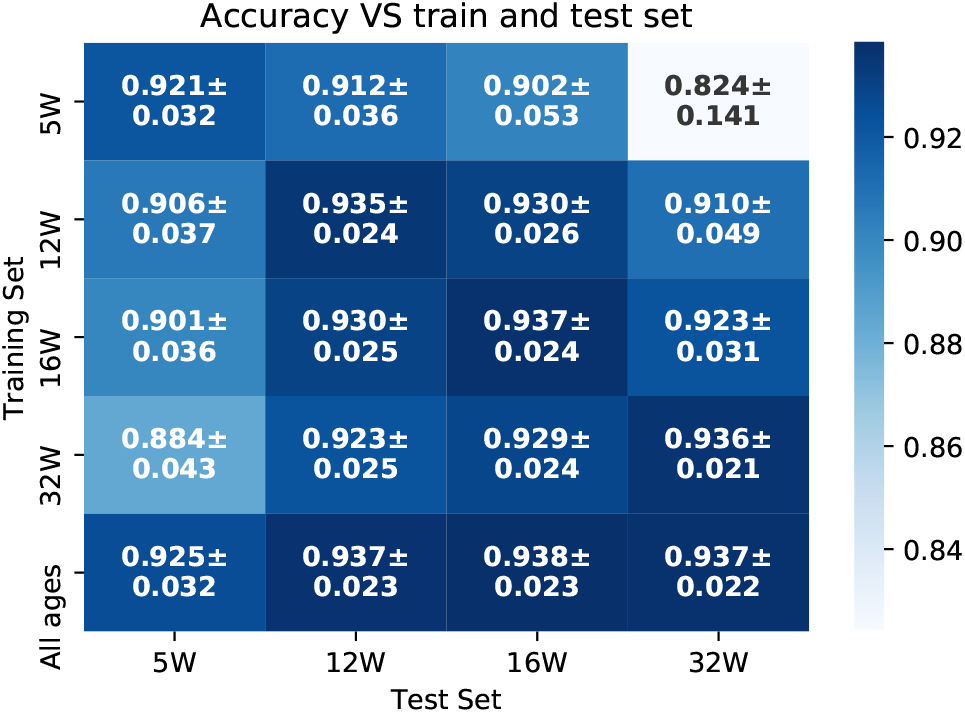
Mean accuracy ± standard deviation for the average accuracy of MU-Net trained and evaluated on different datasets according to mouse age. Networks exclusively trained on older animals achieved lower accuracy when attempting to generalize to the youngest animals, and vice-versa.

### 2.6 Evaluation with a large test dataset

The MU-Net model trained on a train and validation dataset was tested on a large test set of 1782 MRI volumes, acquired from 817 mice with ages ranging from 4 to 60 weeks, and including both WT and HT mice. As the 5-fold cross-validation experiment produced five different MU-Net models, the segmentation maps for the test set were obtained by averaging the prediction maps produced by each of the five models.

Out of the entire test set, segmentation failed completely on two volumes, where no brain mask was detected at all. The remaining 1780 volumes were successfully segmented with an average Dice score of 0.978 ± 0.012 for the brain mask, 0.906 ± 0.041 for the striati, and 0.937 ± 0.035 for the cortex, distributed as illustrated in Fig. 7. There was no significant difference between the segmentation accuracy of male and female animals (*p* > 0.1, unpaired). However, there was a significant difference in accuracy between HT and WT mice (*p* < 0.00001, unpaired) for all ROIs. Dice scores of WT animals were 0.4% higher for the brain mask, 1.7% higher for the cortex, and 1.9% higher for the striati. Applying N3 bias correction on all volumes before segmentation did not result in a significant Dice score difference. A detailed list of the Dice scores for each animal and each ROI is available as an supplementary csv-file.

A visual inspection of the segmentation maps (Fig. 6) revealed that ROIs were qualitatively similar to those obtained on the validation set and displayed in Fig. 2. We observed, however, a visible decrease in performance in the presence of strong ringing artifacts (Fig. 6.b)

**Fig 6.**
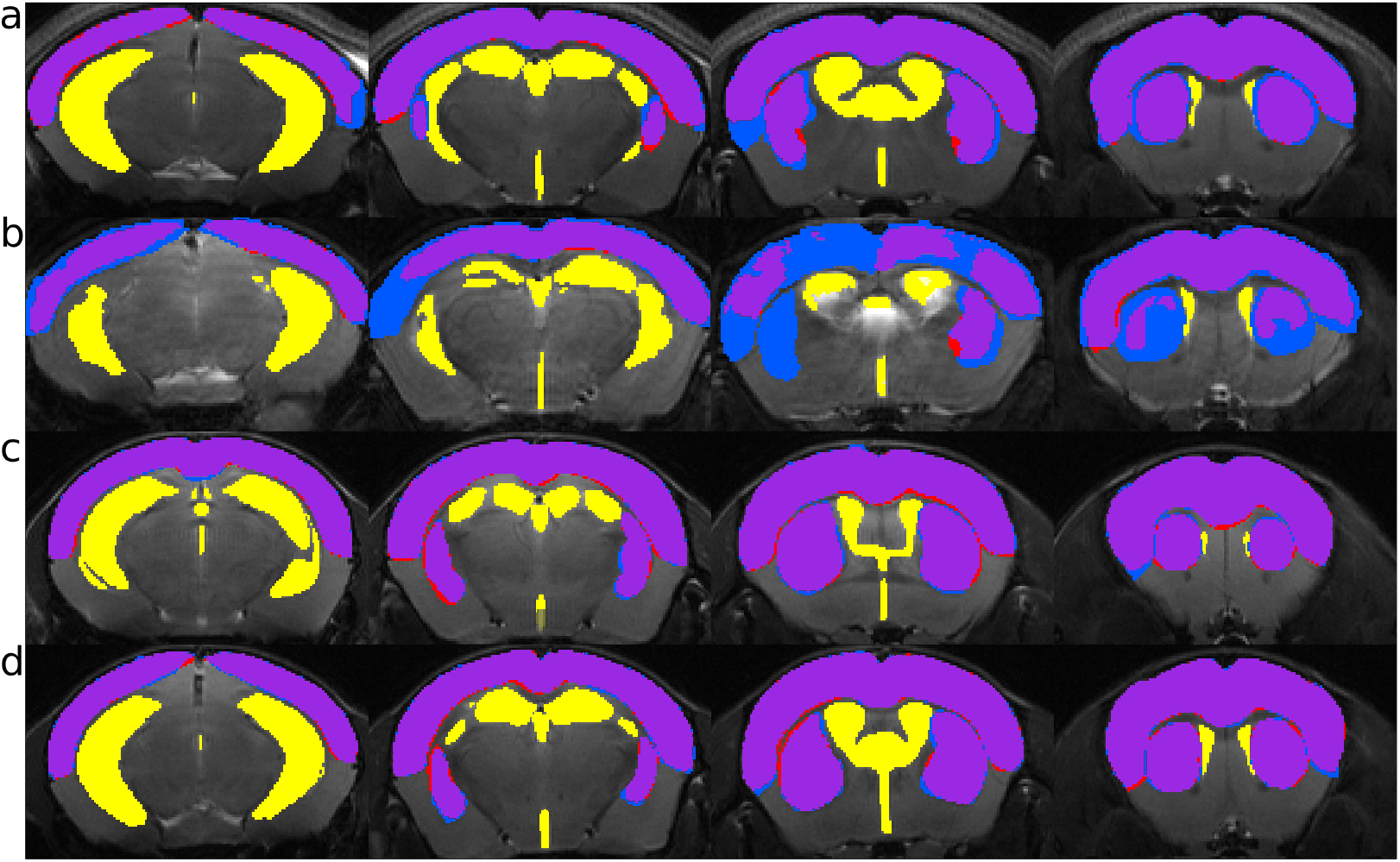
MU-Net segmentation compared to the manual segmentation in four slices of four volumes of the test set. Blue and red indicate, respectively, ground truth and inferred segmentation, purple their overlap (striati and cortex); yellow ROIs (ventricles and hippocampi) are inferred ROIs for which manual annotations were not available. Rows indicate (a) the highest performing volume (mean Dice 0.964, 8 weeks old R6/2 mouse); (b) the lowest performing volume (mean Dice 0.685, 12 weeks old R6/2 mouse); (c) the volume displaying performance closest to the mean performance on the entire test set (Dice 0.923, 12 weeks old Q175DN mouse); (d) one randomly selected volume (Dice 0.919, 8 weeks old Q175DN mouse)

## 3 Discussion

We have presented a multi task deep neural network, MU-Net, for the simultaneous skull-stripping and segmentation of mouse brain MRI. We selected the best performing network among a number of architectures and found it to achieve better segmentation accuracy on the validation set compared to state-of-the-art multi-atlas segmentation procedures, with a markedly lower segmentation time. We then evaluated the performance of MU-Net on a large and heterogeneous test set of 1,782 mice from 10 different studies of Huntington disease, with varying ages and genetic backgrounds (WT as well as HT Q175 and R6/2 variants). In this test set, we measured average Dice scores of 0.978, 0.906 and 0.937 for the brain mask, striati and cortex, rivaling human-level performance. We additionally trained MU-Net for the segmentation of high resolution mouse MRIs of the MRM Neat atlas into 37 regions of interest measuring an average Dice score of 0.820. Hence, we argue that the employment of deep neural networks for the segmentation of animal MRI is a promising strategy for the reduction of both rater bias and segmentation time.

To put the Dice scores in context, Dice scores between two human experts have ranged from 0.80 to 0.90, depending on ROI, for mouse brain MRI segmentation [6]. For different segmentation tasks in brain MRI in general, including human data, inter- and intra-rater Dice score have ranged between 0.75 to 0.96 [6–8]. The Dice scores of MU-Net exceeded these Dice between two human experts, suggesting human-level performance. In addition, the Dice score of MU-Net for skull-stripping was higher than Dice score from the skull-stripping CNN implemented by Roy et al. (0.949) [21]. Obviously, comparing previously reported Dice scores to our segmentation accuracy measures must be done with care as these vary across different studies, segmentation tasks, and datasets, and the confounding factors include image resolution, presence of artifacts and noise, rater expertise, and the choice of ROIs.

While Roy et al. [21] proposed a CNN for skull-stripping for mouse MRI, to our knowledge this work represents the first CNN performing both region segmentation and skull-stripping in mouse brain MRI. The advantages of CNNs to the atlas-based region segmentation [9, 14, 15] are clear. First, compared to atlas-based segmentation MU-Net is much faster and produces accurate results without pre-processing. Second, we found MU-Net to be significantly more accurate than the state-of-the-art STEPS multi-atlas segmentation [15] on anisotropic, relatively quick to acquire MR images favored in pre-clinical drug and biomarker discovery applications. Third, we found MU-Net to perform better than or equally well compared to STEPS on isotropic, high-resolution MR images with relatively long acquisition times, favored in basic research.

We observed that the accuracy of atlas-based methods can vary markedly based on the specific use case depending on the number of outlined ROIs, voxel-size, and image quality. The best performance was achieved using advanced atlas-based methods [15] on the high resolution data [33] with a densely labeled atlas of 37 ROIs, and the lowest using a majority voting rule on a sparsely outlined atlas with a low resolution along the fronto-caudal direction. Our results indicate that under ideal conditions multi-atlas segmentation can achieve accuracy comparable with MU-Net, with a drawback of a long segmentation time. However, where multi-atlas methods underperform, MU-Net still achieved high levels of accuracy, comparable with the Dice score between different manual segmentations.

Interestingly, MU-Nets trained on automatic STEPS multi-atlas segmentations achieved higher Dice score with the ground truth than STEPS, highlighting the generalization ability of MU-Net. This supports the use of atlas based segmentation methods to augment MRI segmentation datasets suggested in [19], leveraging unlabeled data. The results obtained by training on STEPS segmentations alone remain, however, of insufficient quality to eliminate the need for manual annotations in the training data, as the CNN attempts to replicate any form of systematic error present in the atlas-based labeling procedure.

In literature both 3D and 2D implementations of CNNs are available for different segmentation tasks [19, 34, 35], and other architectural variants have been proposed: Roy et al. [19] added dense connections [36] in the convolution blocks of U-Net while keeping the number of output channels constant; Han and Ye [30] proposed two variants based on signal processing arguments for the reduction of artifacts in a sparse image reconstruction task. We, however, found that a more complex model did not improve and in fact lowered the accuracy of our results, perhaps given the simplicity of the task. Thus, in agreement with Isensee et al. [37], we found that a 2D approach was preferable to 3D approach in the presence of anisotropic voxels. We also found the Dice loss to be sufficient to effectively train our model without the addition of a cross-entropy loss.

Much like the human eye, MU-Net was not significantly affected by the presence of the bias field, and did not benefit from N3 bias correction. Correcting for the bias field might still be beneficial as it depends on the specific experimental setup, and thus N3 bias correction might avoid specializing the network to one particular acquisition procedure. For this reason, we release the trained parameters of the model for MU-Net trained on both the non-corrected and the N3-corrected data.

To ensure the network generalizes to a wide age range, our results indicate that the distinctive features present before adulthood need to be adequately represented in the training data. This is evidenced by the degraded performance observed when testing networks trained on 5-week old mice on the volumes acquired from older ones, and vice-versa. As mice are typically weaned at 3-4 weeks and attain sexual maturity at 8-12 weeks [38], 5-week old mice are not adults. In contrast, training solely on male mice did not significantly influence MU-Net’s performance on female animals.

An obvious limitation of our approach is its specialization for the specific MRI contrast the algorithm is trained on, in this case T_2_ MRI. Expanding our approach to a different contrast would require expanding the training set, image translation or transfer learning. Even considering our limitations, MU-Net successfully generalized to a variety of transgenic mice in an age range wider than that of the training set.

The employment of CNNs for the segmentation of mouse brain MRI provides a number of benefits for preclinical researchers. Beyond allowing for the employment of large datasets in a time-efficient manner, the ability to generalize and abstract from the training data results in more robust and reproducible predictions. We can thus expect these methods to reduce the confounding effect of intra- and inter-rater variability inherent in manual segmentation procedures while streamlining animal MRI experimental pipelines.

## 4 Methods

### 4.1 Primary Data

#### 4.1.1 Animals

A total of 849 mice (Charles River Laboratories, Germany) were used: 32 mice for the train and validation set and 817 mice for the test set. Train and validation set animals were scanned at four different ages (5 weeks, 12 weeks, 16 weeks, 32 weeks) resulting in 128 volumes, and did not include any HT genotypes. All train and validation set animals were males.

The test set animals were part of 10 studies scanned at a single or multiple ages from 4 up to 60 weeks, and included both WT and several HT genotypes: R6/2, Q175, Q175DN, Q111, Q50 and Q20 (Supplementary Table S1), for a total of 1,782 MRI scans. The groups included both males and females. These volumes were acquired as part of ten studies of Huntington’s disease, kindly provided by the CHDI ‘Cure Huntington’s Disease Initiative’ foundation.

All mice were housed in groups of up to 4 per cage (single sex) in a temperature (22±1°C) and humidity (30-70%) controlled environment with a normal light-dark cycle (7:00-20:00). All animal experiments were carried out according to the United States National Institute of Health (NIH) guidelines for the care and use of laboratory animals, 336 and approved by the National Animal Experiment Board. 337

#### 4.1.2 MRI

Mice were anesthetized using isoflurane (5% for induction, 1.5-2% maintenance) in 70%/30% mix of N_2_/O_2_ carrying gas, fixed to a head holder and positioned in the magnet bore in a standard orientation relative to gradient coils. Respiration rate and temperature were monitored using PC-SAMS software and Model 1030 Monitoring & Gating System, Small Animal Instruments, Inc., Stony Brook, NY. The temperature was maintained at ~ 37 °C using Small Animal Instruments feedback water heating system.

All acquisitions were performed using a horizontal 11.7 T magnet with a bore size of 160 mm, equipped with a gradient set capable of maximum gradient strength of 750 mT/m and interfaced to a Bruker Avance III console (Bruker Biospin GmbH, Ettlingen, Germany). A volume coil (Bruker Biospin GmbH, Ettlingen, Germany) was used for transmission and a surface phased array coil for receiving (Rapid Biomedical GmbH, Rimpar, Germany). T_2_ weighted anatomical images were acquired using a TurboRARE sequence with effective TR/TE = 2500/36 ms, 8 echoes, 12 ms inter-echo distance, matrix size 256×256, FOV 20.0×20.0 mm2, 31 0.6 mm thick coronal slices, −0.15 mm interslice gap, and 8 averages. Concerning the test data, MRI experimental parameters only differed in acquiring 19 0.7 mm thick contiguous coronal slices.

Volumes within each study were manually segmented by an experienced rater, who had received a training and passed the qualification tests according to SOP (Standard Operating Procedure) for volumetric analysis in mice. Different studies were analyzed by different raters. Each training volume was manually segmented by drawing the brain mask and delineating 4 regions of interest: cortex, hippocampi, striati and ventricles. The brain mask did not include the olfactory bulb or the cerebellum. For the test set, only 3 regions were manually labeled: brain mask, cortex and striati.

### 4.2 MRM NeAt dataset

The MRM NeAt dataset includes atlases of 10 individual T_2_*-weighted in vivo brain MR images of 12-14 weeks old C57BL/6J mice; each with 37 labelled anatomical structures (listed in Figure 4) in addition to the brain mask [33]. This dataset was downloaded from https://github.com/dancebean/mouse-brain-atlas, where an improved atlas is available (bias correction has been applied, left and right labels have been separated and 4th ventricle label added). This dataset was used to evaluate the STEPS algorithm by Ma et al. [15] and is used here for the purpose of comparing MU-Net and STEPS on a larger number of ROIs on isotropic resolution MRI. As detailed in [33], T_2_-weighted MR data with a voxel-size of 0.1 mm^3^ requiring about 2.8 hours of scan time were acquired with a 3D large flip angle spin echo sequence using a super-conducting 9.4T/210 mm horizontal bore magnet (Magnex) controlled by an ADVANCE console (Bruker) and equipped with an actively shielded 11.6 cm gradient set (Bruker, Billerica, MA).

### 4.3 Validation and metrics

To assess the overlap between the ground truth and the predicted segmentation masks, we used the Dice coefficient [29]. The Dice coefficient is defined as two times the size of the intersection over the sum of the sizes of the two regions:

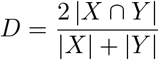

This coefficient ranges from 0, meaning no overlap, to 1, indicating a complete overlap between the two regions. A visual rendition for the intuition behind the Dice score is provided in the supplementary Figure S1.

Each experiment reported in sections 2.2 to 2.3 was validated by splitting the training and validation dataset according to a 5-fold cross validation scheme. Unless otherwise specified, training and validation data included animals of all ages. Volumes were distributed in each fold according to the individual identity of each animal, preventing the use of the volumes from the validation animals for training. The animals were randomly assigned to each fold once, and the same animals remained assigned to their respective folds through all experiments. This resulted in a validation set of 6 or 7 animals and a training set of 25 or 26 mice. These correspond to an identical number of MRI volumes when considering each animal at one specific timepoint only, and to four times as many volumes (24-28 validation volumes, 100-104 training volumes) in all other cases. For the Ma et al. [15] data, 5-fold cross validation results in 8 volumes used for training or as registration atlases, and 2 for testing.

Unless otherwise specified, we used a paired permutation test to evaluate the significance of differences between the Dice scores obtained by different methods, pairing the results obtained on the same MRI volumes. The unpaired permutation test was used instead when comparing results obtained on different volumes, for example, when comparing the accuracy of a model on volumes from younger mice with that of the same model on older mice. We performed permutation tests using 100,000 iterations, and considered differences in average to be significant when *p* < 0.05.

### 4.4 Neural Networks

We segmented mouse brain MRI volumes into five regions (cortex, hippocampi, ventricles, striati and background) simultaneously generating brain mask binary segmentation via a multi-task learning approach. We refer to the first task as the region segmentation and to the latter task as skull-stripping. We implemented and compared different variants of the U-Net architecture.

#### 4.4.1 Architectures

The MU-Net architecture (Fig. 1) is composed of one encoding path and one decoding path. The encoding path includes 4 stages, each articulated in one dense block followed by a 2×2 max-pooling layer. The last feature map feeds into the bottleneck layer, a 64 channel 5×5 convolutional layer with batch normalization [39] connecting the deepest layer of the encoding path with the decoding path.

The decoding path is composed of 4 more blocks alternating one un-pooling layer [40] and one dense block. Un-pooling operations effectively replace up-convolution layers in U-Net without any learnable parameters, while preserving spatial information. These layers operate by simply placing the elements of the un-pooled layer in the position of the respective maximum activation from the corresponding pooling operation, and setting the rest to zero. Skip connections concatenate the output of each dense layer in the encoding path with the respective un-pooled feature map of the same size before feeding it as input to the decoding dense block.

The output of the last decoding layer acts as the input of two different classification layers, which share the same feature representation up to this point: a 1×1 single channel convolution with a sigmoid activation function, and a 1×1 5 channels layer followed by a softmax activation function, for the skull-stripping task and the region classification task, respectively.

##### Dense block

Each dense block includes 3 convolutional layers preceded by leaky ReLU activation [41] layers and batch normalization. In the models including dense connections [36] the input of each convolution is composed by concatenating the outputs of the previous convolutions within the same block and the block’s input. In models where these dense connections are not employed each LeakyReLU / BatchNormalization /convolution sequence simply feeds into the next.

All 3 convolutions are padded and result in 64 output channels, in analogy with Roy et al. [19]. The first and second convolutions employ 5×5 filters, while the third employs a 1×1 filter. This becomes especially relevant in the presence of dense connections, acting as a bottleneck for the 64×3 channels of the concatenated inputs and compressing the size of the feature maps.

##### Dual Framing connections

Dual framing connections refer to additional skip connections proposed by Han and Ye [30] in the Dual Frame U-Net model. This architecture was proposed for computed tomography reconstruction from sparse data based on signal processing arguments to reduce artifacts and improve recovery of high frequency edges. We investigated whether it could prove advantageous in our segmentation tasks. Dual framing connections consist in the subtraction of the input of each dense block on the encoding path from the output of the respective dense block of the same size on the decoding path, and as such the implementation of these connections does not increase the number of model parameters.

##### 3D implementation

A 3D implementation should in principle provide better results by taking into account the features of the adjacent slices, whereas a 2D networks evaluates each coronal slice independently. However, the larger number of parameters also increases the risk of overfitting, and the lower resolution in the anterior-posterior axis compared to the in-plane resolution might constitute confounding factors in the presence of 3D pooling operations.

For these reasons, we confronted 2D and 3D implementations of our network, using 5×5×5 filters and 2×2×2 max-pooling layers, replacing the filters and pooling layers described above. This results in 16008076 and 10286344 parameters for the 3D networks with and without in-block skip connections, respectively. Corresponding 2D networks contain 3297676 and 2087944 parameters, respectively. Thus, opting for a 3D architecture increases the number of parameters by factors of 4.85 and 4.93.

#### 4.4.2 Loss function

Recent literature suggests that Dice-based loss functions [19, 28, 34] would constitute an improvement over cross-entropy losses for the segmentation of medical images [42]. We optimized a joint loss function *L*, that is the sum of two Dice loss functions corresponding to the the skull-stripping (*L_SS_*) and the region classification task (*L_RS_*). Let *p*(*i*) be the predicted probability of voxel *i* of belonging to the brain mask, and *g*(*i*) the ground truth for voxel *i* (*g*(*i*) = 1 if the voxel is in the brain mask). Further, let *p_l_*(*i*) and *g_l_*(*i*) be the same quantities for label *l* (*l* = 1, 2, 3, 4, 5 referring to cortex, hippocampi, ventricles, striati, background) encoding the ground truth as a one-hot vector. Then, the loss function can be written as:

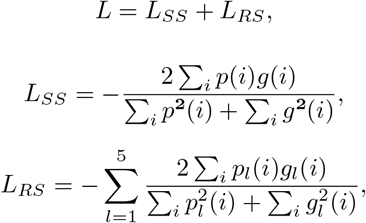

In this formulation each task (skull striping and region segmentation) is defined independently from the other.

#### 4.4.3 Training

The networks were implemented using the PyTorch framework and trained with stochastic gradient descent using Adam optimizer [27] with the default parameters (initial learning rate of 0.001, *β*_1_ = 0.9, *β*_2_ = 0.999 and no weight decay) on an NVIDIA GeForce GTX 1080 GPU. Each training cycle required approximately 12 hours. Training and validation datasets were built according to the 5-fold cross validation scheme described in section 4.3.

Data augmentation was applied on training data. It included random rescaling between 1.01 and 0.95 and random rotation of ±5°. Smaller sizes were preferred in rescaling to decrease memory requirements. Each transformation was applied with a 50% probability. For the same memory reason, a bounding box was created for each volume using the annotated brain mask as a reference. Each volume was individually normalized to 0 mean and unit variance.

Networks used for the comparison on the data released by Ma et al. [15] were trained for up to 24 hours on one Nvidia Volta V100 each, provided by the CSC - IT Center for Science, Finland.

#### 4.4.4 Auxiliary bounding-box network

As MU-Net was trained after cropping the volumes to a bounding box, we trained a lighter 2D network to run a first estimate for the brain mask at inference time from the complete volume. This auxiliary network follows exactly the same architecture of MU-Net, omitting any framing or dense connections, and limiting the number of channels to 4, 8, 16 and 32, from the shallowest to the deepest layer. This results in a network with a total amount of 122454 trained parameters, used to automatically draw a bounding box and run the much heavier MU-Net on the volume thus identified without any manual intervention.

### 4.5 STEPS multi-atlas segmentation

Before registration, each volume underwent non-parametric N3 bias field correction [31] implemented within the ANTS toolset [43]. Taking each volume as reference, all other volumes were then registered with an affine transformation using FSL FLIRT [44] using a correlation ratio cost function, and then nonlinearly registered via FSL FNIRT [45, 46] with the aid of the manually drawn brain mask. Label fusion is achieved with the STEPS algorithm distributed in the NiftySeg package [16, 32].

STEPS depends on the number of templates employed and the standard deviation of its Gaussian kernel. We performed a grid search to select the optimal parameters, randomly selecting 10 volumes and labeling them using STEPS. We sampled the standard deviation of the Gaussian kernels between 0.5 and 6 with a stride of 0.5, and the number of templates ranged between 1 and 20 randomly selected volumes. This same process has been performed both using diffeomorphic registration and using affine registration only (supplementary Figures 2 and 3), selecting 16 templates and kernel standard deviation of 1.5 for the diffeomorphic case, and 18 templates with kernel standard deviation of 2.5 for the affinely registered volumes. Exploring both grids required in total 287 hours.

Each volume was then segmented using these parameters, randomly selecting an appropriate number of mice as templates for the STEPS algorithm as emerged from the parameter grid search outlined above. We repeated this procedure randomly selecting the same number of templates from mice of the same age only. The mice randomly selected as reference atlases were selected from the training set associated to each volume according to the same 5-fold cross validation scheme used to train the CNNs as outlined in section 4.3.

When evaluating STEPS on MRM NeAt dataset, we used scripts provided by Ma et al. [15] at https://github.com/dancebean/multi-atlas-segmentation as this implementation is optimized using this dataset.

### 4.6 Post-processing

The only post-processing steps applied on the segmentation maps were the filling of holes in the resulting 3D volume, the selection of the largest connected component as the brain mask for the skull-stripping task, and assigning all voxels predicted as non-brain to the background class.

**Fig 7.**
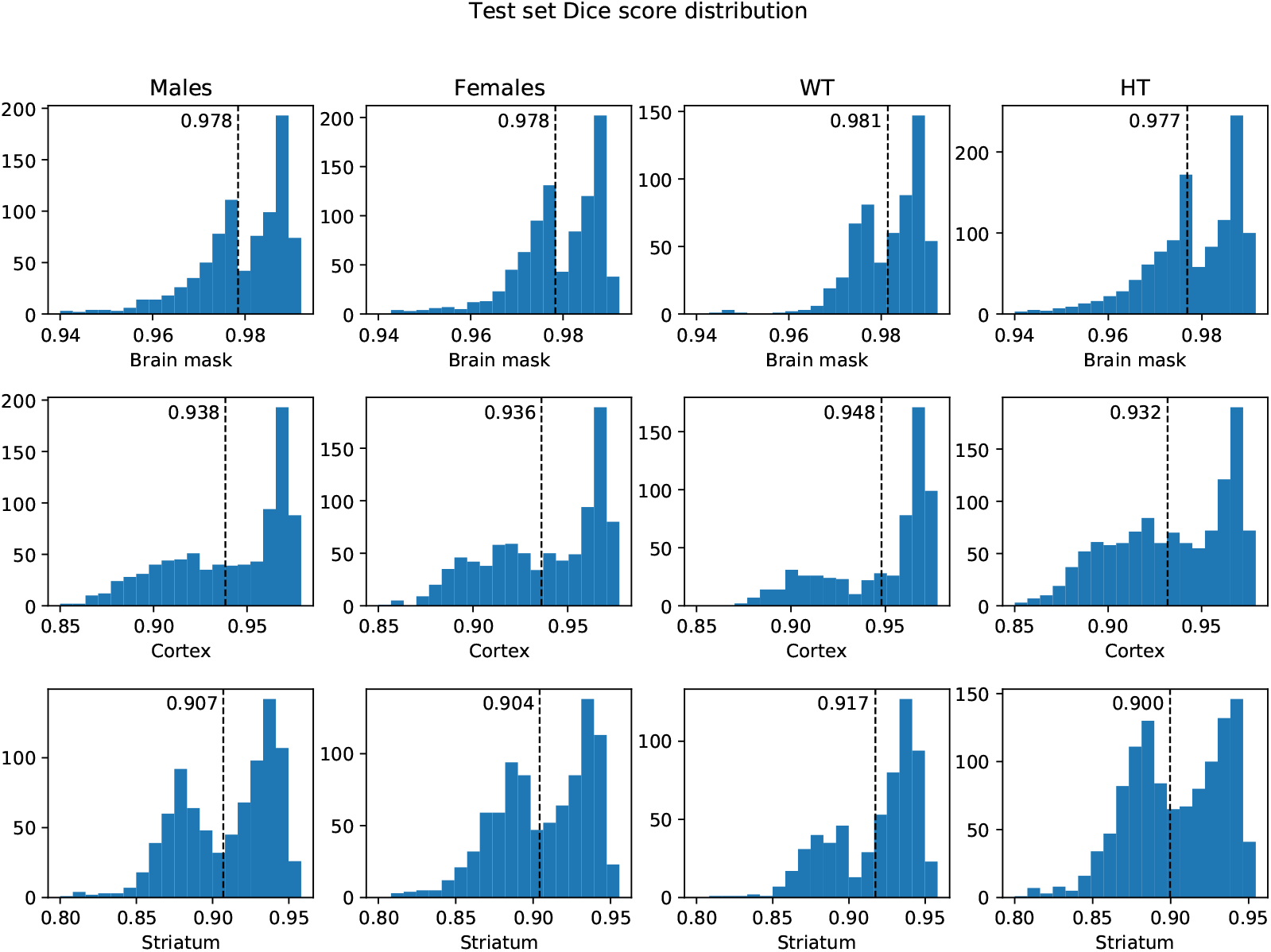
Test set Dice score distribution for the brain mask, cortex and striati ROIs. Males and Females include all mice of each gender, both WT and TG. Likewise, WT and TG include both males and females.

## 5 Declarations

### 5.1 Data availability statement

MU-Net code and trained model are freely available at https://github.com/Hierakonpolis/MU-Net. A tutorial of usage of MU-Net is available at https://github.com/Hierakonpolis/NN4Kubiac The training and validation dataset is property of Charles River Discovery Services, and the test dataset is property of CHDI ‘Cure Huntington’s Disease Initiative’ foundation. The MRM NeAt dataset is freely available at https://github.com/dancebean/mouse-brain-atlas. All the Dice scores between MU-Net and manual segmentations are available as supplementary files to this manuscript.

### 5.2 Contribution statement

J.T., O.G., and F.G. conceived the study, R.D.F. A.Si, and J.T. designed the study, R.D.F. designed the MU-Net network and wrote the software, R.D.F. and A.Si analyzed the data, J.M.V. contributed to the methods, A.Sh. assembled the datasets, R.D.F. and J.T. wrote the paper with substantial input from all the other authors. All authors commented on the manuscript.

## 5.3 Acknowledgments

R.D.F.’s work has received funding from the European Union’s Horizon 2020 Framework Programme under the Marie Skłodowska Curie grant agreement No #691110 (MICROBRADAM) and J.M.V.’ work was founded from Marie Skłodowska Curie grant agreement No #740264 (GENOMMED). The content is solely the responsibility of the authors and does not necessarily represent the official views of the European commission.

The authors wish to acknowledge CSC – IT Center for Science, Finland, for computational resources.

We also extend our thanks to the Academy of Finland, grants (#275453 to A.S. and #298007 to O.G. #316258 to J.T.) and to the CHDI ‘Cure Huntington’s Disease Initiative’ foundation, for kindly providing us with the test data employed in this work. We acknowledge a grant S21770 from European Social Fund to J.T.

## References

1. Cunha L, Horvath I, Ferreira S, Lemos J, Costa P, Vieira D, et al. Preclinical imaging: an essential ally in modern biosciences. Molecular diagnosis & therapy. 2014;18(2):153–173.

2. Matthews PM, Coatney R, Alsaid H, Jucker B, Ashworth S, Parker C, et al. Technologies: preclinical imaging for drug development. Drug Discovery Today: Technologies. 2013;10(3):e343–e350.

3. Febo M, Foster TC. Preclinical magnetic resonance imaging and spectroscopy studies of memory, aging, and cognitive decline. Frontiers in aging neuroscience. 2016;8:158.

4. Anderson RJ, Cook JJ, Delpratt N, Nouls JC, Gu B, McNamara JO, et al. Small animal multivariate brain analysis (SAMBA)-a high throughput pipeline with a validation framework. Neuroinformatics. 2019;17(3):451–472.

5. Calabrese E, Badea A, Cofer G, Qi Y, Johnson GA. A diffusion MRI tractography connectome of the mouse brain and comparison with neuronal tracer data. Cerebral cortex. 2015;25(11):4628–4637.

6. Ali AA, Dale AM, Badea A, Johnson GA. Automated segmentation of neuroanatomical structures in multispectral MR microscopy of the mouse brain. Neuroimage. 2005;27(2):425–435.

7. Yushkevich PA, Piven J, Hazlett HC, Smith RG, Ho S, Gee JC, et al. User-guided 3D active contour segmentation of anatomical structures: significantly improved efficiency and reliability. Neuroimage. 2006;31(3):1116–1128.

8. Entis JJ, Doerga P, Barrett LF, Dickerson BC. A reliable protocol for the manual segmentation of the human amygdala and its subregions using ultra-high resolution MRI. Neuroimage. 2012;60(2):1226–1235.

9. De Feo R, Giove F. Towards an efficient segmentation of small rodents brain: a short critical review. Journal of neuroscience methods. 2019;.

10. Pagani M, Damiano M, Galbusera A, Tsaftaris SA, Gozzi A. Semi-automated registration-based anatomical labelling, voxel based morphometry and cortical thickness mapping of the mouse brain. Journal of neuroscience methods. 2016;267:62–73.

11. Schwarz AJ, Danckaert A, Reese T, Gozzi A, Paxinos G, Watson C, et al. A stereotaxic MRI template set for the rat brain with tissue class distribution maps and co-registered anatomical atlas: application to pharmacological MRI. Neuroimage. 2006;32(2):538–550.

12. Sharief AA, Badea A, Dale AM, Johnson GA. Automated segmentation of the actively stained mouse brain using multi-spectral MR microscopy. Neuroimage. 2008;39(1):136–145.

13. Lerch JP, Sled JG, Henkelman RM. MRI phenotyping of genetically altered mice. In: Magnetic Resonance Neuroimaging. Springer; 2011. p. 349–361.

14. Bai J, Trinh TLH, Chuang KH, Qiu A. Atlas-based automatic mouse brain image segmentation revisited: model complexity vs. image registration. Magnetic resonance imaging. 2012;30(6):789–798.

15. Ma D, Cardoso MJ, Modat M, Powell N, Wells J, Holmes H, et al. Automatic structural parcellation of mouse brain MRI using multi-atlas label fusion. PloS one. 2014;9(1):e86576.

16. Cardoso MJ, Leung K, Modat M, Keihaninejad S, Cash D, Barnes J, et al. STEPS: Similarity and Truth Estimation for Propagated Segmentations and its application to hippocampal segmentation and brain parcelation. Medical image analysis. 2013;17(6):671–684.

17. LeCun Y, Bengio Y, Hinton G. Deep learning. nature. 2015;521(7553):436.

18. Wachinger C, Reuter M, Klein T. DeepNAT: Deep convolutional neural network for segmenting neuroanatomy. NeuroImage. 2018;170:434–445.

19. Roy AG, Conjeti S, Navab N, Wachinger C. QuickNAT: Segmenting MRI Neuroanatomy in 20 seconds. arXiv preprint arXiv:180104161. 2018;.

20. Ronneberger O, Fischer P, Brox T. U-net: Convolutional networks for biomedical image segmentation. In: International Conference on Medical image computing and computer-assisted intervention. Springer; 2015. p. 234–241.

21. Roy S, Knutsen A, Korotcov A, Bosomtwi A, Dardzinski B, Butman JA, et al. A 6 deep learning framework for brain extraction in humans and animals with traumatic brain injury. In: Biomedical Imaging (ISBI 2018), 2018 IEEE 15th International Symposium on. IEEE; 2018. p. 687–691.

22. Szegedy C, Liu W, Jia Y, Sermanet P, Reed S, Anguelov D, et al. Going deeper with convolutions. In: Proceedings of the IEEE conference on computer vision and pattern recognition; 2015. p. 1–9.

23. Chou N, Wu J, Bingren JB, Qiu A, Chuang KH. Robust automatic rodent brain extraction using 3-D pulse-coupled neural networks (PCNN). IEEE Transactions on Image Processing. 2011;20(9):2554–2564.

24. Oguz I, Zhang H, Rumple A, Sonka M. RATS: rapid automatic tissue segmentation in rodent brain MRI. Journal of neuroscience methods. 2014;221:175–182.

25. Xie L, Qi Y, Subashi E, Liao G, Miller-DeGraff L, Jetten AM, et al. 4D MRI of polycystic kidneys from rapamycin-treated Glis3-deficient mice. NMR in Biomedicine. 2015;28(5):546–554.

26. Valverde JM, Shatillo A, De Feo R, Gröhn O, Sierra A, Tohka J. Automatic rodent brain mri lesion segmentation with fully convolutional networks. In: International Workshop on Machine Learning in Medical Imaging. Springer; 2019. 3 p. 195–202.

27. Kingma DP, Ba J. Adam: A method for stochastic optimization. arXiv preprint arXiv:14126980. 2014;.

28. Sudre CH, Li W, Vercauteren T, Ourselin S, Cardoso MJ. Generalised dice overlap as a deep learning loss function for highly unbalanced segmentations. In: Deep learning in medical image analysis and multimodal learning for clinical decision support. Springer; 2017. p. 240–248.

29. Dice LR. Measures of the amount of ecologic association between species. Ecology. 1945;26(3):297–302.

30. Han Y, Ye JC. Framing U-Net via deep convolutional framelets: Application to sparse-view CT. IEEE transactions on medical imaging. 2018;37(6):1418–1429.

31. Sled JG, Zijdenbos AP, Evans AC. A nonparametric method for automatic correction of intensity nonuniformity in MRI data. IEEE transactions on medical imaging. 1998;17(1):87–97.

32. Cardoso MJ, Modat M, Ourselin S, Keihaninejad S, Cash D. STEPS: multi-label similarity and truth estimation for propagated segmentations. In: Mathematical Methods in Biomedical Image Analysis (MMBIA), 2012 IEEE Workshop on. IEEE; 2012. p. 153–158.

33. Ma Y, Smith D, Hof PR, Foerster B, Hamilton S, Blackband SJ, et al. In vivo 3D digital atlas database of the adult C57BL/6J mouse brain by magnetic resonance microscopy. Frontiers in neuroanatomy. 2008;2:1.

34. Milletari F, Navab N, Ahmadi SA. V-net: Fully convolutional neural networks for volumetric medical image segmentation. In: 2016 Fourth International Conference on 3D Vision (3DV). IEEE; 2016. p. 565–571.

35. Çiçek Ö, Abdulkadir A, Lienkamp SS, Brox T, Ronneberger O. 3D U-Net: learning dense volumetric segmentation from sparse annotation. In: International conference on medical image computing and computer-assisted intervention. Springer; 2016. p. 424–432.

36. Huang G, Liu Z, Van Der Maaten L, Weinberger KQ. Densely connected convolutional networks. In: Proceedings of the IEEE conference on computer vision and pattern recognition; 2017. p. 4700–4708.

37. Isensee F, Petersen J, Klein A, Zimmerer D, Jaeger PF, Kohl S, et al. nnu-net: Self-adapting framework for u-net-based medical image segmentation. arXiv preprint arXiv:180910486. 2018;.

38. Dutta S, Sengupta P. Men and mice: relating their ages. Life sciences. 2016;152:244–248.

39. Ioffe S, Szegedy C. Batch normalization: Accelerating deep network training by reducing internal covariate shift. arXiv preprint arXiv:150203167. 2015;.

40. Noh H, Hong S, Han B. Learning deconvolution network for semantic segmentation. In: Proceedings of the IEEE international conference on computer vision; 2015. p. 1520–1528.

41. Maas AL, Hannun AY, Ng AY. Rectifier nonlinearities improve neural network acoustic models. In: Proc. icml. vol. 30; 2013. p. 3.

42. Karimi D, Salcudean SE. Reducing the Hausdorff Distance in Medical Image Segmentation with Convolutional Neural Networks. arXiv preprint arXiv:190410030. 2019;.

43. Avants BB, Tustison N, Song G. Advanced normalization tools (ANTS). Insight j. 2009;2:1–35.

44. Jenkinson M, Smith S. A global optimisation method for robust affine registration of brain images. Medical image analysis. 2001;5(2):143–156.

45. Jenkinson M, Beckmann CF, Behrens TE, Woolrich MW, Smith SM. Fsl. Neuroimage. 2012;62(2):782–790.

46. Andersson JL, Jenkinson M, Smith S, et al. Non-linear registration aka Spatial normalisation FMRIB Technial Report TR07JA2. FMRIB Analysis Group of the University of Oxford. 2007;.

