## Supplementary figures and images for "Automated joint skull-stripping and segmentation with Multi-Task U-Net in large mouse brain MRI databases"

### Supplementary figure 1

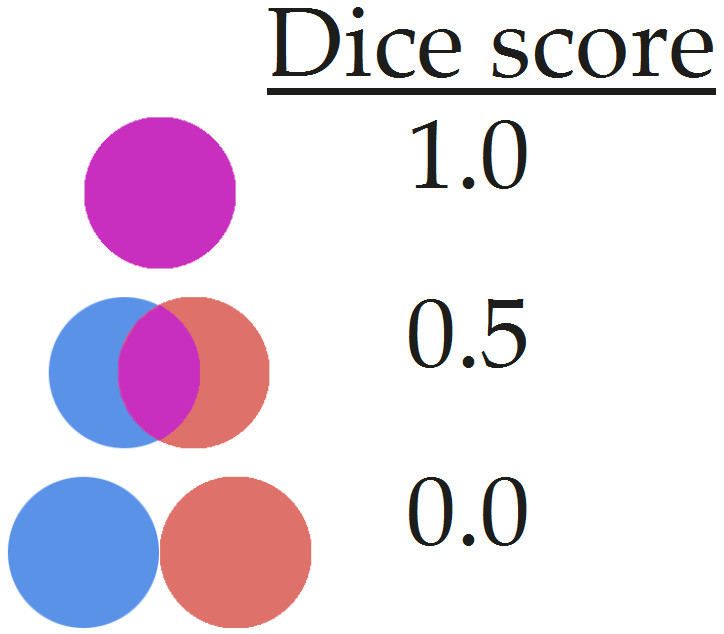

### Supplementary figure 2

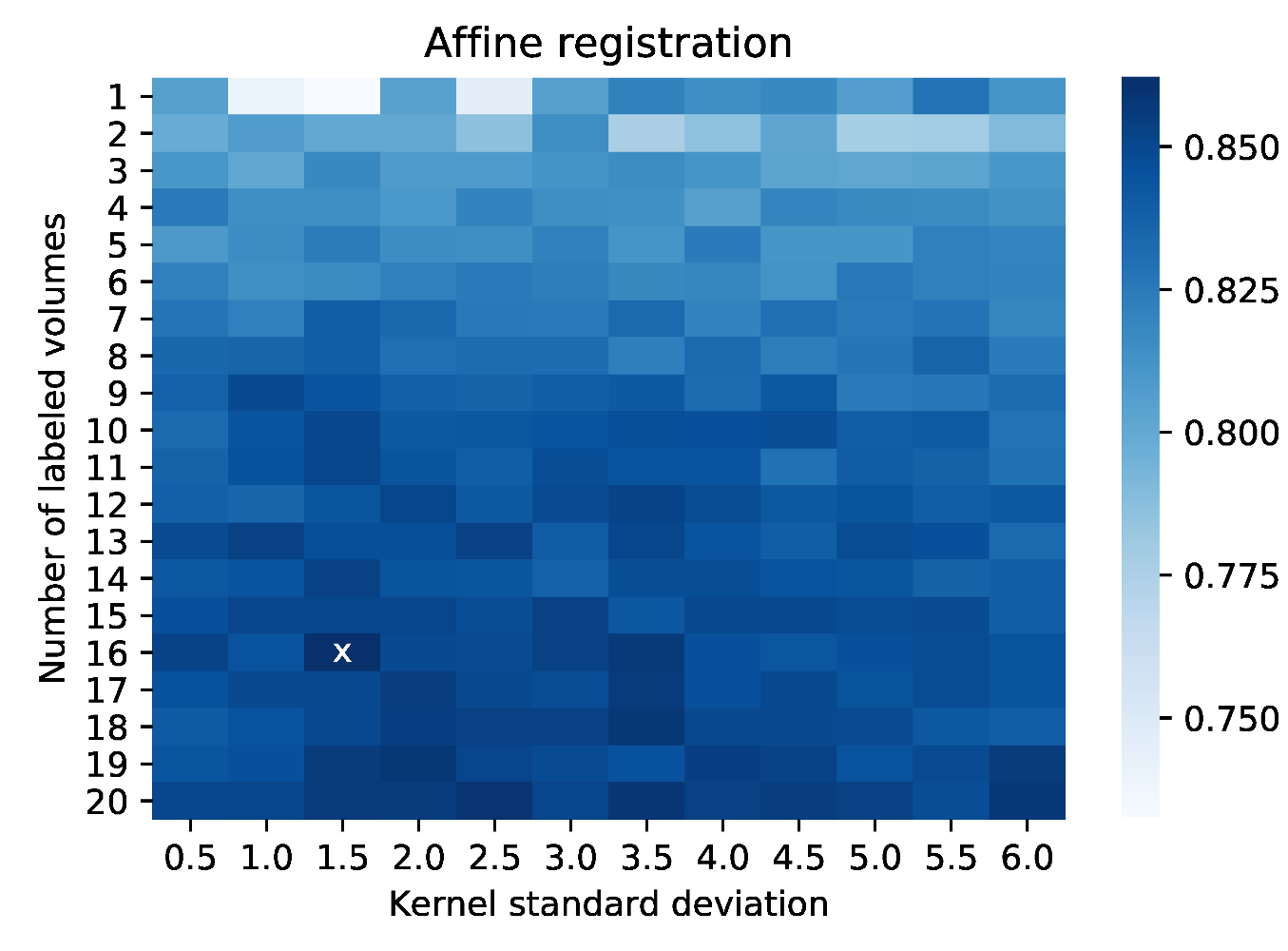

### Supplementary figure 3

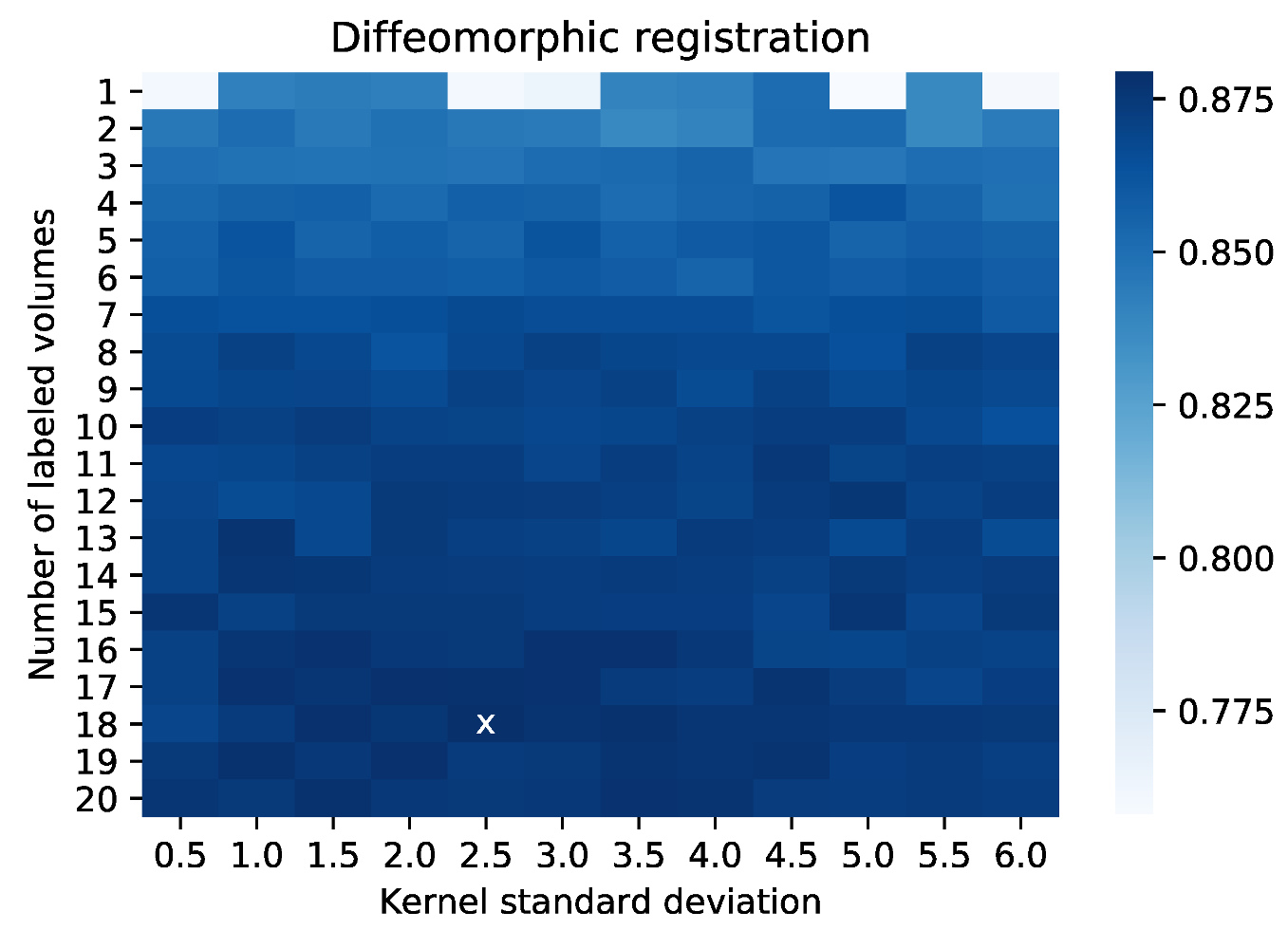
